# Accumulation of genetic variants associated with immunity in the selective breeding of broilers

**DOI:** 10.1101/763656

**Authors:** Angela Zou, Kerry Nadeau, Pauline W. Wang, Jee Yeon Lee, David S. Guttman, Shayan Sharif, Doug Korver, John H. Brumell, John Parkinson

## Abstract

To satisfy an increasing demand for dietary protein, the poultry industry has employed genetic selection to increase the growth rate of broilers by over 400% in the past 50 years. Although modern broilers reach a marketable weight of ∼2 kg in a short span of 35 days, a speed twice as fast as a broiler 50 years ago, the expedited growth has been associated with several negative detrimental consequences. Aside from heart and musculoskeletal problems, which are direct consequences of additional weight, the immune response is also thought to be altered in modern broilers. Given that identifying the underlying genetic basis responsible for a less sensitive innate immune response would be economically beneficial for poultry breeding, we decided to compare the genomes of two unselected meat control strains that are representative of broilers from 1957 and 1978, and a current commercial broiler line. Through analysis of genetic variants, we developed a custom prioritization strategy to identify genes and pathways that have accumulated genetic changes and are biologically relevant to immune response and growth performance. Our results highlight two genes, TLR3 and PLIN3, with genetic variants that are predicted to enhance growth performance at the expense of immune function. Placing these new genomes in the context of other chicken lines, reveal genetic changes that have specifically arisen in selective breeding programs that were implemented in the last 50 years.

## INTRODUCTION

Poultry production is key to meeting a rising global demand for high quality dietary protein. In efforts to meet this demand, the poultry industry has employed intensive breeding programs aimed at selecting for lines that promote growth and enhance the efficiency of feed conversion ratios[1]. In the last 50 years, such selection has resulted in broilers displaying remarkable gains in growth and efficiency in feed conversion, such that the modern broiler attains a marketable weight in less than half the time, using half the amount of feed relative to breeds produced in the 1950’s. However, such gains can be associated with compromised health; selection for rapid growth without due regard to physiological support systems can lead to musculoskeletal disorders and sudden heart failures, and greater susceptibility to disease compared to older breeds[2, 3]. Of particular concern to the broiler industry is necrotic enteritis (NE), caused by the bacterial pathogen *Clostridium perfringens*. Global losses due to NE are estimated to be $2B/year and they are expected to rise[4].

To help mitigate against losses due to NE, the industry has relied on the use of sub-therapeutic quantities of antibiotic (antibiotic growth promotants, AGPs) that are added to chicken feed to protect from NE and enhance production efficiency[5]. Unfortunately, with the global emergence of antimicrobial resistance, many countries are seeking to ban the use of AGPs and the incidence of enteric diseases in chicken is expected to rise[6, 7]. Consequently, with selective breeding predicted to maximize growth potential in the next decade, the industry has shifted towards maintaining poultry health to minimize economic losses. To help support these programs, there is a need to identify the genetic modifications that have resulted in declined health and disease susceptibility. In attempts to meet this need, several groups have employed genomic approaches to characterize a diverse set of breeds, including native Taiwanese chickens[8], experimental lines that serve as animal models[9], as well as domestic layer and broiler lines[10, 11]. Aside from identifying genes that dictate physical characteristics such as plumage or skin color, these studies have identified numerous selective sweeps[8, 10, 12], reflecting regions with reduced heterozygosity as a consequence of positive selection. Such sweeps have revealed the fixation of alleles predicted to be beneficial. For example, specific alleles of the gene encoding thyroid stimulating hormone receptor (TSHR), which is critical for proper metabolic regulation and reproduction, have been shown to be fixed in all modern chickens, but not in the genomes of the Red Jungle Fowl[10, 13, 14] or chickens dating from ∼280 B.C. to ∼1700 A.D[13]. Beyond allelic variation, several studies have also focused on the role of copy number variation on disease and phenotype diversity [11, 15, 16]. For example, a study of 12 diversified chicken genomes identified a number of copy number variable regions covering genes that include *FZD6* and *LIMS1*, which have been associated with increased resistance to Marek’s disease[11], while a comparison of Leghorn and Fayoumi chickens identified a duplication of BD7, involved in host innate immune response, in the latter [15]. Typically, these studies have focused on pairwise comparisons of individual lines, as opposed to genetic variation that occurs within the context of a breeding program. The availability of lines with shared ancestry that have undergone selective breeding over a number of decades, offers an opportunity to explore how the accumulation of genetic traits might impact phenotype.

In this study, we performed a comprehensive genomic analysis of domestic chicken breeds, focusing on broilers that underwent intensive selective breeding in the last 50 years. We generated genome sequence data for broiler lines generated in 1957, 1978, and 2016, which offer, for the first time, snapshots of genetic changes introduced during the breeding program, as well as insight into the underlying genetics of accelerated growth and changes in immune function. Additionally, we integrate whole genome datasets of other domestic chicken breeds and performed single nucleotide polymorphism calling on all data sets. We have since then identified genetic variants unique and common to different breeds that we suspect may contribute to breed-specific phenotypes. More importantly, we are able to identify genes and pathways involved in metabolism and innate immune responses that have been altered by artificial selection, with some pathways showing potential in modifying modern broiler responses to foreign pathogens,that represent novel avenues for future investigations focusing on improving poultry health.

## RESULTS

### Genome sequencing reveals the modern broiler to be closely related to the 1957 and 1978 lines

The goal of this study was to identify the genetic basis for traits selected for in modern breeding programs, such as growth and feed efficiency, with the purpose of informing on the selection of otherwise unanticipated traits such as skeletal defects, metabolic disorders and altered immune function[2, 3, 17]. To this end we sequenced the genomes of two University of Alberta Meat Control strains, unselected since 1957 and 1978 and representing synthetic crosses of a large number of contemporary commercial broiler strains (designated 1957 and 1978, respectively), and a 2016 commercial Ross 308 broiler (designated Ross308). After DNA extraction, Illumina sequencing resulted in the generation of 950, 956, and 750 million 150 bp reads for 1957, 1978, and Ross 308 respectively, representing from 76X to 92X coverage – ensuring sufficient coverage to accurately infer single nucleotide polymorphisms (SNPs; **Table 1**)[18]. To supplement our analyses, we included genome data from 12 additional breeds (**Supplemental Table 1**) obtained from the National Center for Biotechnology Information Sequence Read Archive (NCBI SRA)[8, 10]. These include: two layers – Swedish white layer (WLA) and commercial White Leghorn (WLB); two broilers – Ross 308 (CB1) and an undisclosed commercial broiler (CB2); two dual purpose chickens bred for both meat and egg production – Rhode Island Red (RIR) and a Taiwan heritage L2 line (L2), a Silkie line (Silkie); three experimental lines – high (High) and low (Low) growth lines established from White Plymouth Rock chickens in 1957[19] and an Obese line (Obese) established in 1955 from a White Leghorn line as a model for autoimmune thyroiditis[20]; and two distinct genome sequences derived from Red Jungle Fowls (RJF and RJFswe).

**Table 1.**
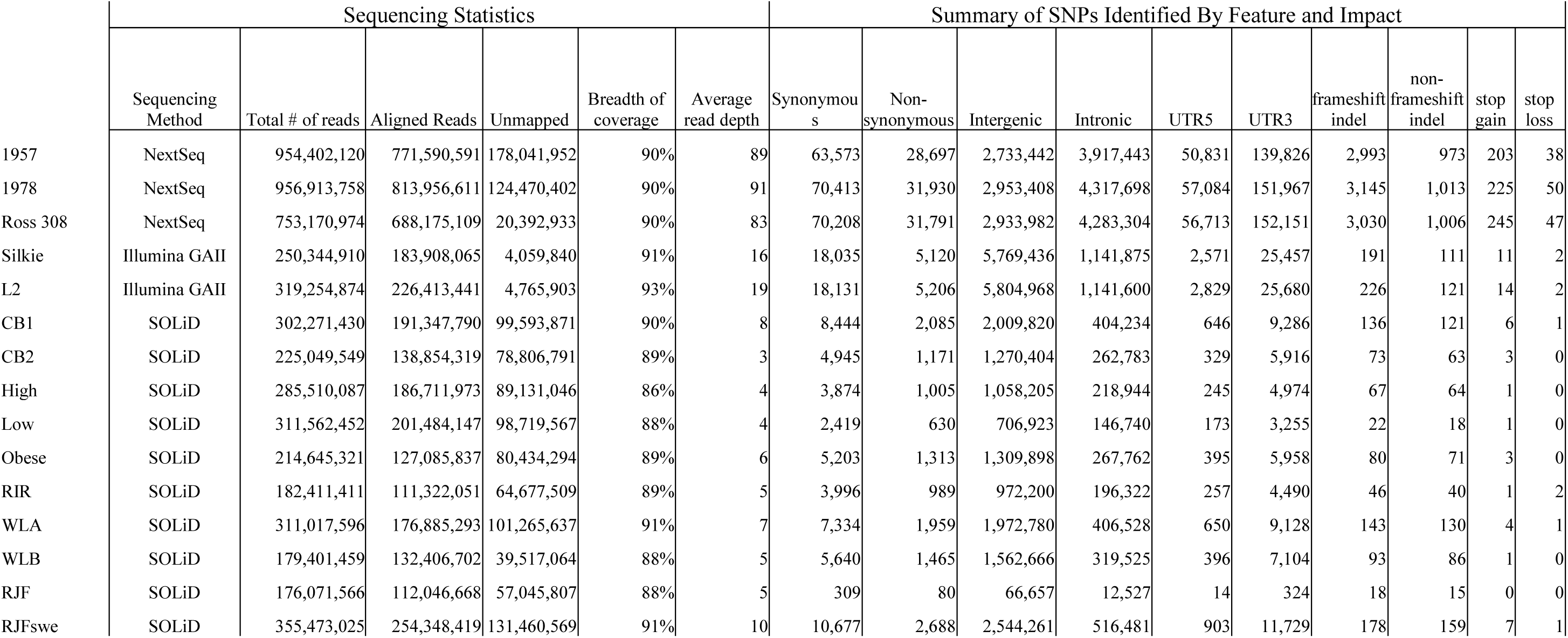
Sequencing and Genomic Variant Statistics for Breeds Used in This Study.

Alignment of sequence reads to the reference chicken genome assembly (GCA_000002315.3)[21] reveal the number of variants to be broadly consistent with known breeding history. For example, the Red Jungle Fowl, aligned against its own genome, had the fewest (80,000) variants. Further, experimental lines (High, Low and Obese) also tended to have fewer variants than the production lines. Lines sequenced with the Illumina platform (1957, 1978, Ross308, Silkie and L2), which yield greater depth of sequence coverage, each had more variant calls than those sequenced with SOLiD technology. Interestingly, unlike other lines, the three lines sequenced here exhibited greater numbers of intronic variants (∼3.9-4.3 million) compared to intergenic variants (∼2.7-2.9 million). Also noteworthy, the second Red Jungle Fowl dataset (RJFswe) exhibited the greatest number of variants amongst those lines sequenced by the SOLiD platform. This increase in variation may reflect both a greater genetic heterogeneity in this line, together with the pooling of DNA from two different sources (8 males were selected from two different zoos in Sweden[10])

To further examine the potential impact of sequencing platform on variant calls, we performed a phylogenetic analysis on the variants common to lines sequenced by SOLiD and Illumina platforms (**Figure 1**). Reassuringly our analysis grouped both Ross 308 datasets (Ross308 and CB1), together with the 1957 and 1978 lines. Furthermore, the clustering of the High and Low lines, as well as the Obese, WLA and WLB lines is consistent with known breeding history[10]. On the other hand our analysis split the CB1 and CB2 lines, which were previously considered close in ancestry. Our analysis also grouped the L2 and the Silkie lines, sequenced by Illumina, with the three broiler lines. The L2 line was established as a closed population from a slow growing Taiwanese line more than 30 years ago and has been selected for both egg and meat production^1^, while the Silkie line, was originally bred for ornamental purposes in the 19^th^ century^2^. Of note, the RJFswe line did not group with the RJF line, suggesting that pooling of samples from two sources may have impacted variant calls in this dataset.

**Figure 1.**
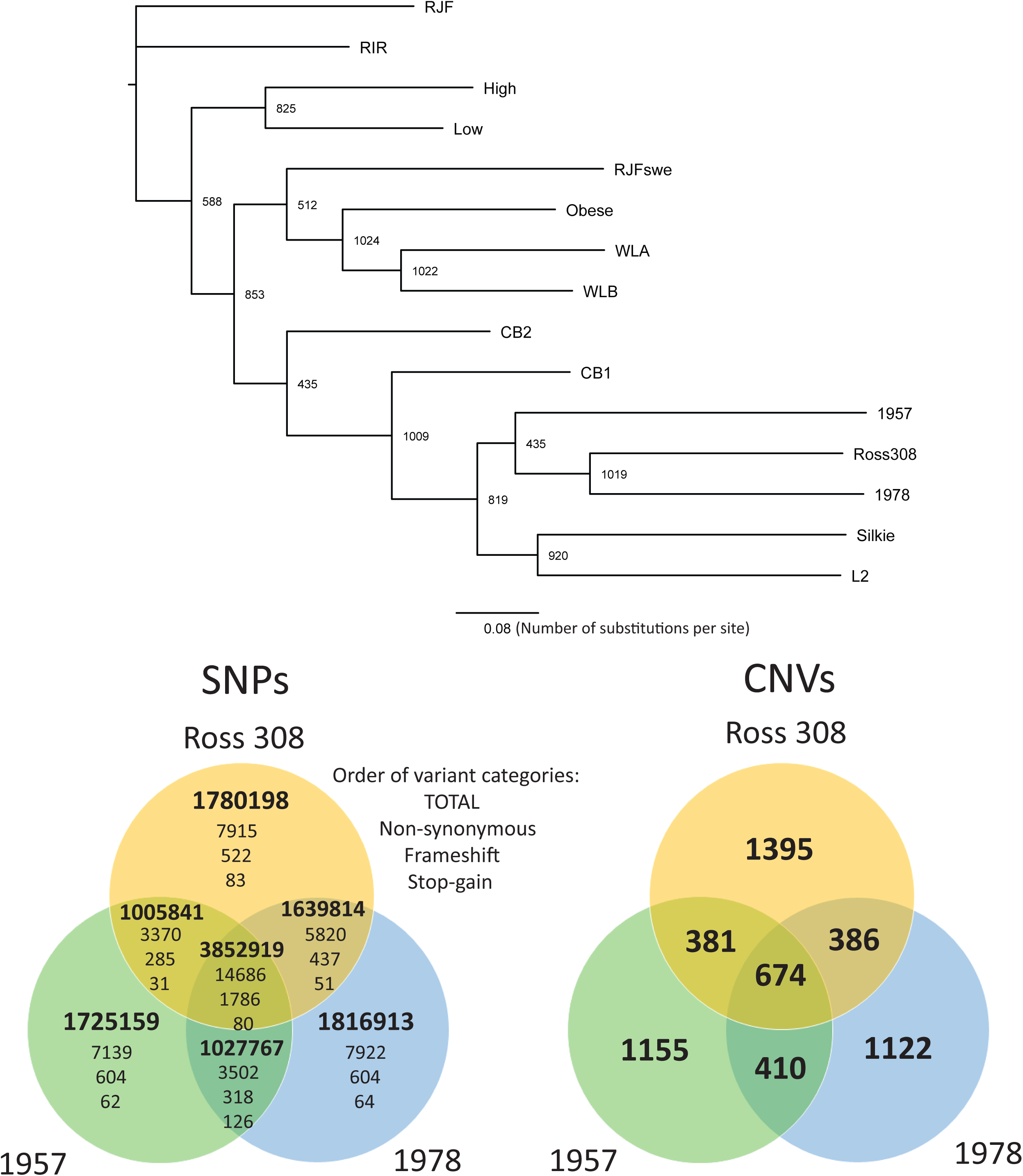
Overview of variant results following SNP and CNV calling. A) Phylogenetic analysis of chicken lines. The phylogenetic tree was reconstructed using PhyML with 1024 bootstrap replicates. Bootstrap values are indicated at the branch points, the scale bar indicates number of nucleotide substitutions per site. Red Jungle Fowl was used to root the tree. Only SNPs identified in at least one Illumina-sequenced and two SOLiD sequenced datasets and occuring within genomic regions covered by more than 5 reads in all SOLiD datasets and more than 10 reads in all Illumina datasets were used in the analysis. (B) and (C) Venn diagrams displaying the number of SNPs (B) and CNVs (C) that are unique and shared between the three genomes sequenced in this study.

Focusing specifically on the three broiler lines (1957, 1978 and Ross308), we found that while each line possessed ∼1.7 – 1.8 million SNPs not present in the other two lines, they share a common core of ∼3.9 million SNPs. Amongst this core were 14,686 non-synonymous SNPs, together with 1796 frameshifts and 80 stop gains – variants that were likely introduced in the common ancestor of the three lines. Further, we found that the 1978 and Ross308 genomes share more SNPs (1,639,814) than the 1957 and the Ross308 genomes (1,005,841). In addition to analyzing single-nucleotide polymorphisms (SNPs), we also compared copy number variants (CNVs; see Methods). CNVs that encompass coding regions have the potential to alter gene expression[22], with a concomitant impact on phenotypic traits. Multiple studies have already identified CNVs in diverse chicken breeds[8, 11, 16, 23-25]. Here we aimed to compare CNVs across our three lines to identify loci with the potential to contribute to phenotypic changes in commercial broilers. Our analyses identified over 2,500 CNVs for each line (**Table 2**), spanning ∼8% of the chicken genome. Of these, 674 are common to all three lines (**Figure 1**), with Ross 308 having the most unique CNVs in terms of number (1395 CNVs) which together account for 30% of the length of all CNV regions found in that line. These regions were associated with copy gains for ∼150-170 genes and copy losses of ∼150-160 genes. Gene set enrichment analysis of CNVs for each set of copy gains or losses revealed no enrichment of any Gene Ontology (GO) terms.

**Table 2.** Summary of CNVs detected in broilers relative to Red Jungle Fowl genome.

|  | **1957** | **1978** | **Ross 308** |
| --- | --- | --- | --- |
| CNVs detected | 2625 | 2587 | 2836 |
| Median CNV size (bp) | 3975 | 3975 | 3900 |
| Average CNV size (bp) | 37696 | 38492 | 34517 |
| Range (min, max) | 1050,<br>9295280 | 1050,<br>8706900 | 1050,<br>9684450 |
| Coverage breadth (%) | 8% | 8% | 8% |
| Duplications | 425 | 512 | 543 |
| Deletions | 2200 | 2075 | 2293 |
| Gains overlapping with genes | 148 | 170 | 168 |
| Losses overlapping with genes | 156 | 159 | 156 |

Notable genes associated with CNVs include two copies of CD8alpha in Ross308, two copies of tumor necrosis factor superfamily member 13b (TNFS13B) in both 1978 and Ross308, and the loss of one copy of Apolipoprotein-AII (APO-AII) in both 1978 and Ross308. CD8alpha is a co-receptor on CD8+ T cells, a specialized group of T-cells that recognizes antigens presented by MHC class I molecules, then respond to damaged cells by triggering production of cytokines and the release of cytotoxic molecules[26]. Related, TNFS13B (tumor necrosis factor superfamily member 13b; alternate name B cell activating factor, BAFF) is a cytokine that binds its corresponding receptors to induce proliferation and differentiation of B cells[27]. Heightened expression of this ligand has been observed in Crohn’s disease patients compared to healthy controls and has been proposed as a potential marker of IBD[28]. Hence, copy number gains in CD8alpha and TNFSF13B in the 1978 and Ross308 lines may be linked to increased inflammation in response to infection. APO-AII, on the other hand, coats lipids to form high-density lipoproteins and promote cholesterol efflux[29, 30]; deficiency in this protein results in high serum cholesterol levels, which has been observed in Ross308 broilers relative to chickens indigenous to Iran[31].

### Enrichment analysis reveals genetic variants associated with specific biological processes that vary across lines

To achieve a broad understanding of the impact of SNPs in each line, a functional enrichment analysis was performed to reveal biological processes significantly enriched in SNPs in the different chicken lines. First, for each gene we define its *SNP density* as the number of mutations divided by the coding sequence length. We then constructed a set of *SNP dense* genes as the 10% of genes displaying the greatest SNP density (and hence most likely impacted by genetic variation) and performed a gene set enrichment analysis for enriched functional categories as defined through GO and pathways defined by the Kyoto Encyclopedia of Genes and Genomes (KEGG; **Figure 2**). Across all lines, 17 GO terms were defined as significantly enriched in SNP dense genes, of these 10 were enriched in more than one line, with the four immune related terms: “Herpes simplex infection” (KEGG: 05168); “Cytokine-cytokine receptor interaction” (KEGG: 04060); “Intestinal immune network for IgA production” (KEGG: 04672); and “Phagosome” (KEGG: 04145), being the most widely represented (significant in 7, 7, 5 and 5 lines respectively). Given the association of these terms with a diverse range of lines (broilers, Silkie, Taiwanese heritage chicken), this enrichment highlights the impact of domestication on the chicken immune response[32, 33]. Not all lines had G.O. terms enriched in SNP dense genes, and while the Ross308 genome sequenced here possessed the most enriched terms (10), only 3 were shared with the previously sequenced genome (CB1), which was additionally enriched in the immune-related term “Defense response to bacterium” (GO:0042742). Of note, Ross308 was uniquely associated with three terms relating to bone morphogenetic proteins (BMPs): “BMP signaling pathway” (GO:0030509); “Cellular response to BMP stimulus” (GO:0071773); and “Response to BMP” (GO:0071772). Such enrichment may reflect changes in pathways contributing to musculoskeletal disorders suffered by modern broilers[34]. For the three broilers that we sequenced, the terms “Defense response to other organism” (GO:0098542) and “Defense response” (GO:0006952) were only enriched in Ross308. Among the genes assigned to these terms are three that potentially link immune response and metabolic stress: TMEM173, a gene that activates type I interferon production and promotes glucose intolerance and tissue inflammation under metabolic stress induced by high fat diets[35]; and the two cathelicidins, CATHB1 and CATH3, antimicrobial peptides that kill bacteria by puncturing their cell membranes. With CATHB1, present in the intestinal tract where it can influence intestinal permeability, potential dysbiosis, and growth performance[36], and CATH3 able to bind lipopolysaccharides and reduce inflammation[37], the association of these categories with Ross308 and not 1957 or 1978, suggest that recent selection may have resulted in compromised immune function. Together these GO enrichment analyses reveal biological processes that may have been altered through breeding programs.

**Figure 2.**
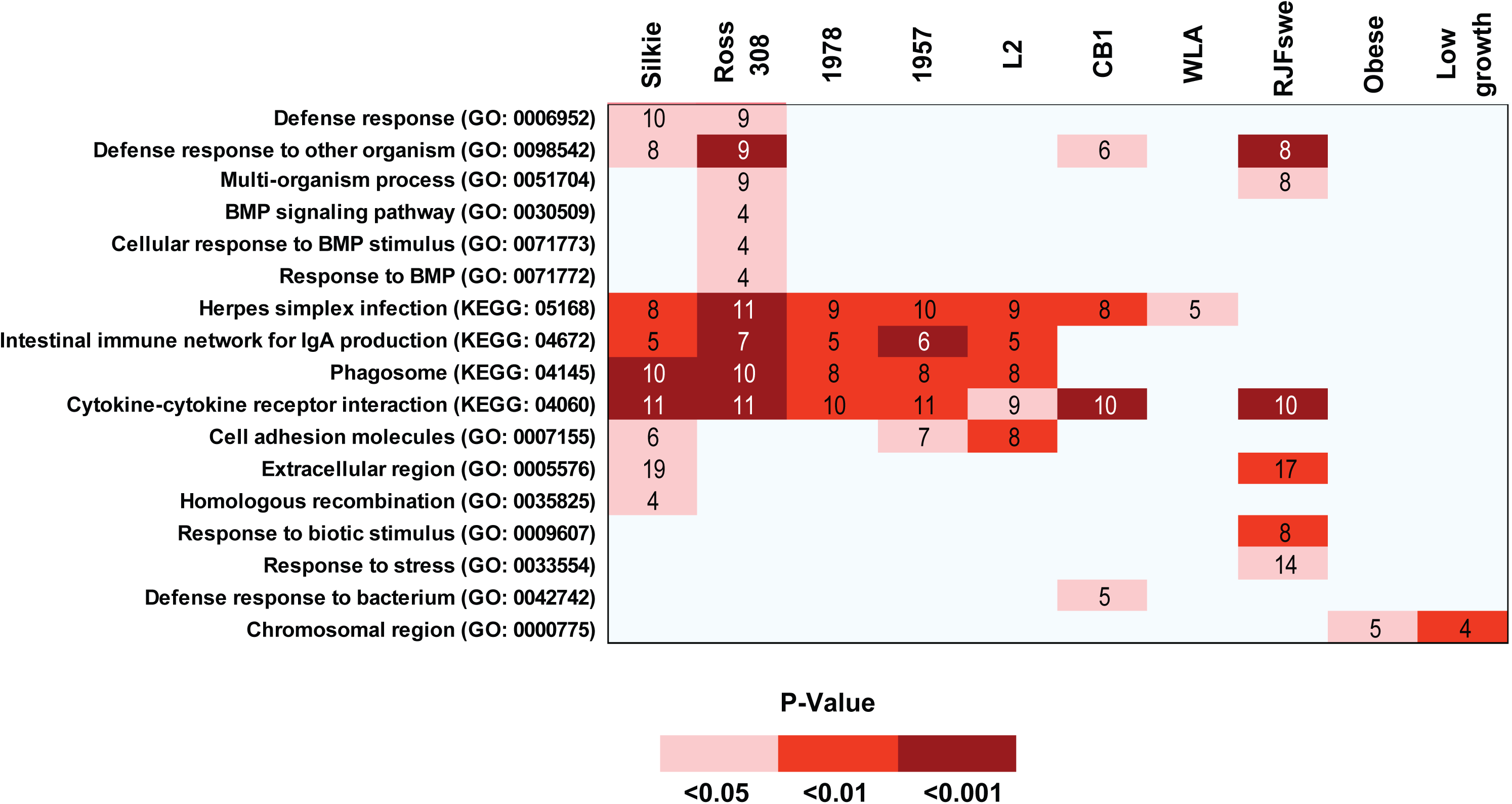
Heat map showing enrichment of GO and KEGG categories of genes containing nonsynonymous SNP densities in the 90th percentile across all breeds. All categories and pathways shown have a FDR-adjusted p-value of less than 0.05 in at least one line.

### Combinatorial prioritization strategy identifies candidate genes for future functional characterization

In the previous section we identified sets of biological processes enriched in SNP dense genes associated with 10 lines; many were associated with growth and immune responses. To identify transcripts that have been functionally impacted in the development of the Ross308 line and lay the framework for future experimental investigations, we developed a novel prioritization scheme (**Figure 3**) based on combining five complementary strategies (see Methods). Note, we refer here to transcripts rather than genes since four of the five strategies rely on normalization of SNPs and SNP effects by transcript length.

**Figure 3.**
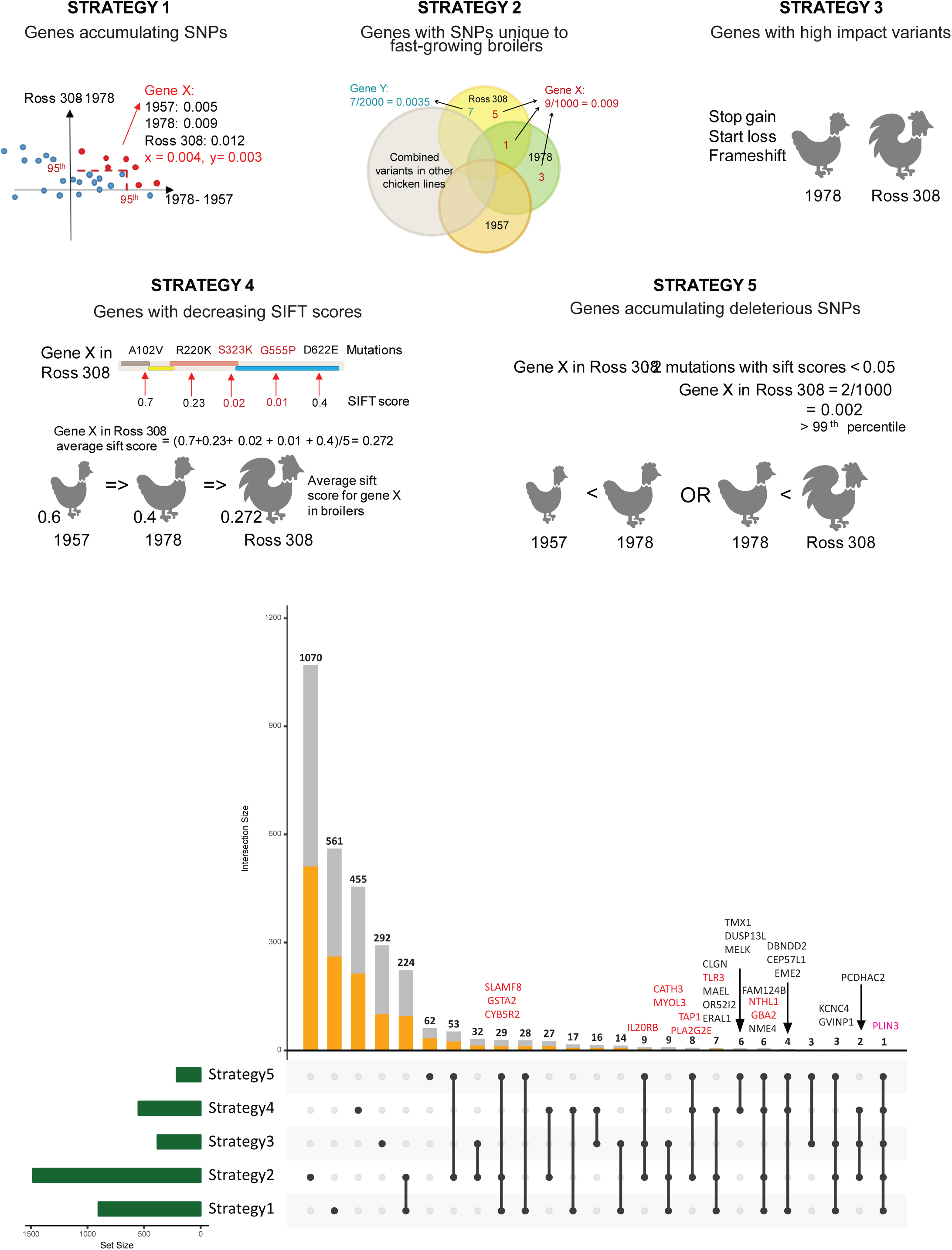
Prioritization strategies used to identify genes associated with selection of fast growing broiler lines. Top panels: Schematic representation of strategies used to prioritize genes (see main text for more explanation. Lower panel, upset plot showing numbers of transcripts identified through each strategy and combination of strategies. Known genes associated with more restricted combinations of strategies are indicated. Genes in red are the 12 manually curated genes mentioned in manuscript.

The first strategy (*Strategy 1*) defines transcripts that have accumulated the greatest number of SNPs over the past 60 years. Here we compared increases in non-synonymous SNP density between the 1957 and the 1978 lines, and the 1978 and Ross308 lines. Under this strategy, we select transcripts that are in the 95^th^ percentile in terms of increase in SNP density either for the 1957/1978 comparison or the 1978/Ross308 comparison. We additionally specify that both changes in SNP density must be positive (i.e. there are more SNPs in more recently bred lines). In *Strategy 2* we define transcripts based on unique SNPs in faster-growing lines. Transcripts are selected based on possessing the highest (90th percentile) density of SNPs that are unique to the 1978 and/or Ross308 lines relative to all other lines. *Strategy 3* focuses on transcripts that possess high impact variants (i.e. frameshift, start loss, and stop gain variants) restricted to either the 1978 or Ross308 lines, but not the 1957 line. Similarly, *Strategy 4* identifies transcripts based on decreasing SIFT scores (a measure of whether a SNP will affect protein function). Under this strategy, a list of transcripts in which the mean SIFT score decreases from the 1957 to 1978 lines and the 1978 to Ross308 lines is first generated. Subsequently, transcripts from this list are selected if they are in the lower 50th percentile of SIFT scores for either the 1978 or Ross308 lines. In the final strategy (*Strategy 5*) we identify transcripts that have accumulated deleterious SNPs. Here we select for transcripts that show an increase in the number of deleterious variants (SNPs with SIFT scores less than 0.05) when comparing the 1957, 1978, and Ross308 broilers and, additionally, possess the highest (99^th^ percentile) density of deleterious variants.

Applying each strategy resulted in the generation of five complementary set of transcripts (**Figure 3**). Strategy 2 resulted in the largest number of transcripts (1477 and strategy 5 the fewest (211). Combining strategies, we find that only 20, of 120 possible combinations of strategies result in unique sets of transcripts. To help prioritize genes for future experimental investigations, we subsequently filtered for transcripts that are identified in at least three strategies (**Figure 3)**. This final filtering step ensures the selection of transcripts with SNPs that result in functional consequences (use of only one or two strategies might result in the selection of transcripts that have simply accumulated many low-impact variants) and identified a total of 78 transcripts. From this list, manual reviews of GO and domain annotations allowed us to identify 12 transcripts with mammalian homologs involved in immune response and/or promoting growth, that have likely been compromised in the selection of the 1978 and Ross308 lines (CATH3, CYB5R4, GBA2, GSTA2, IL20RB, MYOL7L2, NTHI1, PLA2G2E, PLIN3, SLAMF8 and TAP1). After manual review of the potential functional impact of their associated SNPs, we identified TLR3 and a putative homolog of mammalian PLIN3 (ENSGALT00000003248), for more focused analysis.

TLR3 is part of the Toll-like receptor family that plays a crucial role in innate immunity by recognizing double stranded viral RNA and promoting anti-viral defenses in other cells [38]. In the Ross308 line, TLR3 has acquired nine more variants than the 1978 line and 16 more than the 1957 line. Several of these variants are unique to the Ross308 (relative to the other lines considered in this study) and occur in conserved regions (**Figure 4**). These variants include N174H and D466H, both found in the leucine-rich surface that is responsible for recognizing and binding double stranded viral RNA and activating innate immunity. When viewed in the context of other mutations that alter charge (D19V, K43Q, R266S, E283G), we hypothesize that the sensitivity of TLR3 to double stranded viral components could be diminished. This decrease in functional efficacy could be physiologically important for growth performance, for example, a mouse knock out of a toll-like receptor (TLR5) has previously been shown to modulate gut microbiome characteristic of obesity and other metabolic disorders[39].

**Figure 4.**
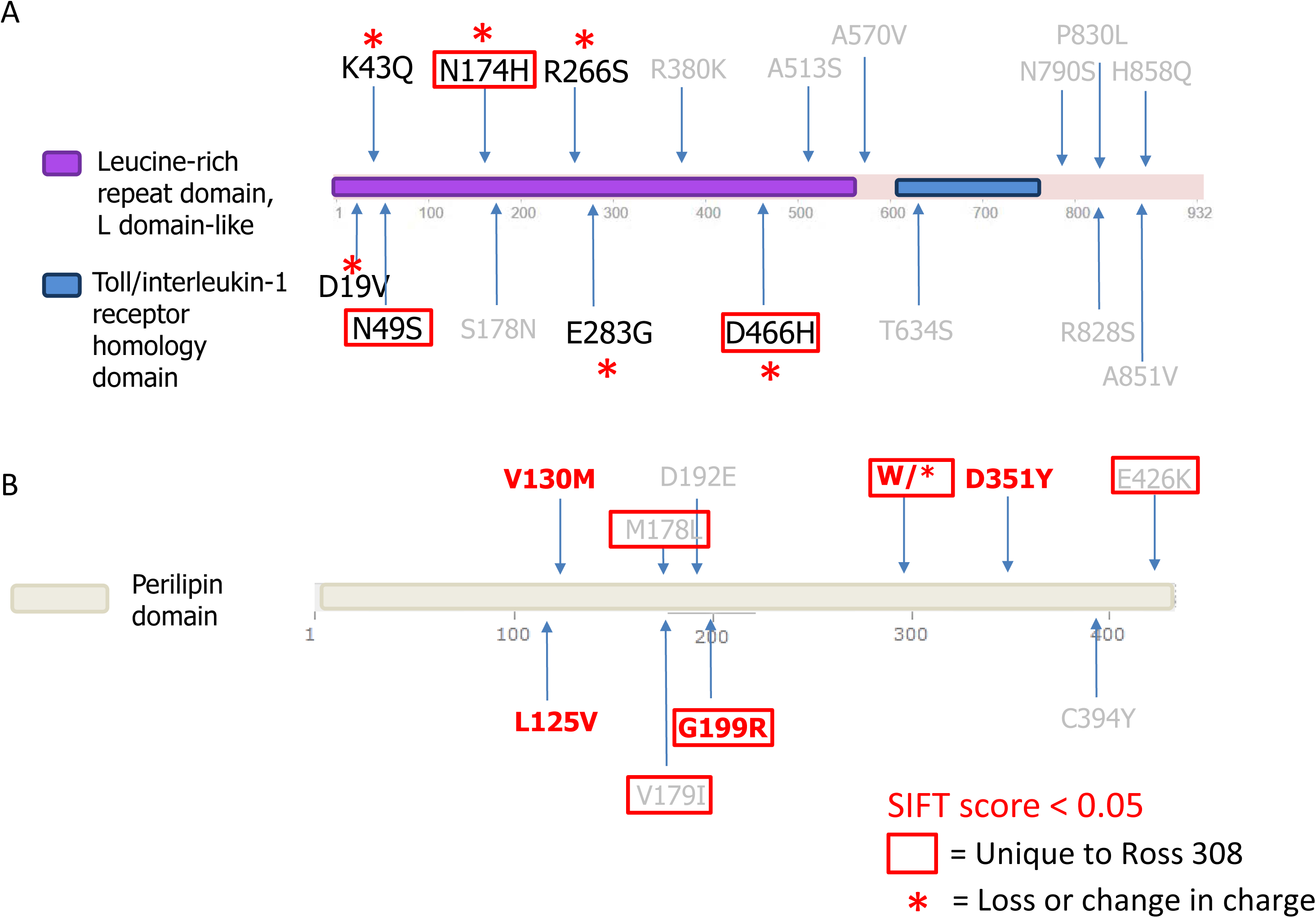
Domain architectures of TLR3 and PLIN3 showing non-synonymous SNPs identified in Ross308. (A) Domain architecture of TLR3, showing leucine-rich domain and the Toll/interleukin-1 receptor homology domain. (B) Domain architecture of PLIN3. Highlighted for both genes are SNPs considered deleterious (SIFT score < 0.05), as well as those that are unique to Ross308 and/or result in a loss or change in charge.

ENSGALT00000003248, a PLIN3 homolog, is the only gene that was identified by all five strategies; of the ten SNPs found in the Ross308 line, five were unique, including the deleterious variant G199R and a stop codon W298X predicted to cause premature termination (**Figure 4**). ENSGALT00000003248 is a gene product of ENSGALG00000002083 and identified by Ensembl as a putative paralog of PLIN3. PLIN3 is a member of the perilipin family, a group of proteins known for regulating lipid metabolism in different cell types. PLIN3 has been observed to respond to stimulation by lipopolysaccharides in neutrophils, while knocking down PLIN3 in neutrophils results in an immediate decrease in lipid droplets biogenesis and prostaglandin E2 production[40], both of which are critical for initiating an appropriate innate inflammatory immune response. Moreover, PLIN3 participates in exercise-induced lipolysis by co-localizing with lipid droplets and interacting with adipose triglyceride lipase[41], while its malfunction can lead to dysregulations in fat oxidation often observed in obese or diabetic patients. Given its role in fat deposition, and that it is the only gene to be prioritized by all five strategies, we hypothesize that alterations in this gene have helped to promote growth in the Ross 308 line.

## DISCUSSION

In this study, we compared genetic variants in the genomes of three broiler lines: two meat control strains representative of broilers in 1957 and 1978, and a 2005 commercial Ross 308 broiler. Our central hypothesis is that immune pathways were compromised during selective breeding targeting increased growth and efficiency. To address this question, we developed a selection strategy to identify genes that have acquired variants over the course of selective breeding programs, impacting their function. Our initial expectation is that this strategy would result in the identification of genes that contribute to both immune and growth-related processes.

Our combinatorial strategy incorporates three considerations neglected by other studies. First, different genes could experience different levels of selective constraints which has been observed in domestication of cattle [42] and maize [43], where variants that may be considered “deleterious” in natural populations are relaxed under changes in environments (for example as might occur in commercial broiler facilities). Second, while previous studies focused on identification of causal variants [44-46], genes can also accumulate multiple mutations which are tolerated individually but may act synergistically to affect gene function. Lastly, this study is relatively unique from a poultry perspective in presenting analyses of genomes that are representative of different time points across commercial broiler genetic selection, offering a novel opportunity to track the accumulation of mutations in genes over time. Consequently, we focused our analyses on longitudinal comparisons of the 1957, 1978, and Ross308 genomes.

Many of the genes that passed our final filter (prioritized by more than 3 strategies) contribute to both immune response and growth (Table 2). TLR3 already has an established role of recognizing viral dsRNA to elicit an innate immune response. However, Toll-like receptors, the superfamily that encompasses TLR3, have also been shown to indirectly influence metabolism[47, 48]. Knocking out TLR2[48] and TLR4[49] can prevent diet-induced obesity and insulin resistance through inflammatory signaling pathways. Conversely, TLR5 knockouts in mice have been shown to induce insulin resistance, hyperphagia, and many other symptoms characteristic of metabolic disorders[39]. Examining the gut microbiome composition in TLR5-deficient mice reveals that it was altered in such a way that it induced low-level inflammatory signaling, this in turn, attenuated insulin signaling to promote insulin resistance and hyperphagia. Similarly, we hypothesize that TLR3, which does not bind fatty acids like TLR2 and TLR4[50, 51], may also mediate metabolism in broiler. Indeed, TLR3 has already been observed to protect mice from colitis due to its ability to sense dsRNA produced by commensal lactic acid bacteria and suppress inflammatory responses reserved for pathogenic bacteria[52]. By measuring dsRNA amounts and inflammatory response following digestion of dsRNA, it was established that dsRNA level in commensal Lactobacillales, which is higher than dsRNA amounts in pathogens such as *Salmonella enterica* serovar *Typhimurium* and *Escherichia coli*, is sufficient to induce anti-inflammatory signals via TLR3. Another study that used a high-density SNP array to investigate polymorphisms in a commercial broiler line also found that TLR3 falls within a region of positive selection[12]. This raises the interesting possibility that mutations that result in decreased TLR3 function may result in poor discrimination between commensal and pathogenic flora of the gut, that while resulting in dysbiosis, may nevertheless improve growth performance.

PLIN3 belongs to the perilipin family of proteins that associate with lipid droplets to regulate lipase access to lipid storage. With known roles in exercise-induced lipid degradation[41] and LPS-induced lipid droplet formation in neutrophils[40], the accumulation of genetic variants in PLIN3 during selective breeding of the Ross308 line suggests that functions may be altered in this line, with potential impact on both lipid accumulation and immune response.

In addition to identifying genes of interest, we observed that all three broilers that we sequenced are over-represented in SNP dense genes from several functional categories such as phagosome, cytokine-cytokine receptor interaction, and intestinal immune network for IgA production. That suggests that commercial broiler immune response, irrespective of lineage, is distinct from the Red Jungle Fowl immune function. The Ross308 line, however, is the only one of the three broilers that we sequenced with SNP dense genes enriched in the GO term: “Defense response to other organisms” (GO:0098542). This result is supported by previous studies; one study found that commercial broilers had a lower febrile response (ability to generate fever) and lower cytokine expression in response to gram-negative bacteria than commercial layers[53]. Comparing the gene expression profiles of two broiler lines and quantifying pathogen burden after infection with *Salmonella* revealed the fast-growing chickens relied on T-cell activation and were slower at clearing *S. enterica* serovar Enteritidis infection than slow growing chickens, which induced macrophage activation post infection[54]. Therefore, from our investigations of enriched G.O. terms (specifically: “Defense response to bacterium” (GO:0042742) and “Defense response to other organisms” (GO:0098542)), these studies support the hypothesis that the Ross308 line may respond differently to pathogenic invasion.

Together this study illustrates how comparisons of the genome sequences derived from temporally related lines can help reveal the genes and biological processes that help drive livestock breeding programs.

## METHODS

### Genome sequencing and data collection of additional datasets

The genomes of two female chickens from each of the University of Alberta Meat Control strains unselected since 1957 and 1978, respectively, and a commercial Ross 308 broiler (Aviagen North America Inc., Huntsville, Alabama) were generated as follows:

Genomic DNA was isolated from 30 ul of chicken blood samples using the Promega Wizard Genomic DNA Purification Kit (Promega, Madison, WI, USA) according to the manufacturer’s instructions. Genomic DNA was quantified with a Qubit 2.0 fluorometer using the Qubit dsDNA BR assay kit (Life Technologies, Carlsbad, CA) and diluted in buffer EB to the 20 ng/ul input for library preparation. DNA samples were also assessed for quality by gel electrophoresis. Library preparation was performed following the Illumina Truseq DNA PCR-Free LT Library Prep Kit (Illumina, San Diego, CA) protocol using 1ug of isolated genomic DNA. The input DNA was sheared to 350bp using Covaris microTUBES and the S2 Ultrasonicator (Covaris, Woburn, MA). End repair, A-tailing, adaptor ligation and library nomalization were performed according to Illumina’s instructions. Libraries were quantified for normalization by qPCR using the KAPA Library Quantification Kit for Illumina sequencing platforms (KAPA Biosystems, Boston, MA). The final libraries were also assessed using the DNA1000 kit for the Bioanalyzer 2100 (Agilent Technologies, Palo Alto, CA). The final libraries were denatured and diluted to a final loading volume of 1.3 ml at a concentration of 1.8pM. Sequencing was performed on the NextSeq500 sequencer (Illumina, San Diega, CA), according to the manufacturer’s instructions, using the 300 cycle High Output V2 sequencing kit and a spike-in of 8% PhiX control library, generating 150 bp paired end reads. Sequence data is available through the sequence repository archive, hosted by the NCBI, under BioProject PRJNA561435.

Additional genomes were downloaded from the sequence read archive (SRA) at the National Center for Biotechnology Information (NCBI)[55] and include the genomes of Taiwan L2 and Silkie chickens (SRP022583), together with ten lines generated through pooled blood samples (SRP022583): a White Leghorn line developed at the Swedish University of Agricultural Sciences (WLA), a commercial White Leghorn line (WLB), an Obese line, Rhode Island Red (RIR), Ross 308 (CB1), a second unidentified commercial broiler line (CB2), high- and low-growth lines, Red Jungle Fowl from a Swedish zoo population (RJFswe), and the partly inbred UCD 001 line used to generate the reference chicken genome (RJF).

### Alignment and variant detection of Illumina data sets

For lines sequenced using the Illumina platform (1957, 1978, and Ross 308 broilers sequenced here, as well as Taiwan L2 and Silkie), adapters and low quality reads were identified and filtered using FastQC[56] and Trimmomatic[57]. Remaining reads were mapped separately to the *Gallus gallus* reference genome (galGal5, January, 2016[21]) with BWA 0.7.5[58] using default settings. Picard tools (http://broadinstitute.github.io/picard) removed duplicates in the alignment. Genome analysis toolkit(GATK)[59] was used for indel re-alignment and base score recalibration, where high confidence variants were first detected with HaplotypeCaller and used to recalibrate base scores. HaplotypeCaller was subsequently used to call SNPs, with the minimum emission confidence threshold set at a phred score of 15 and the minimum calling confidence threshold set at a phred score of 50. Detected variants were further filtered with Variant Filtration (parameters used were QUAL / DP < 2.0, FS > 60.0, MQ < 40.0, MQRankSum < −12.5, ReadPosRankSum < −12.5).

Putative copy number variants (CNVs) of the 1957, 1978, and Ross 308 genomes were detected using CNVnator[60] (bin size set to 45) and further filtered by size (> 1000 bp) and reads mapping location (fraction of reads that can map to 2 or more locations must be less than 0.5). When comparing CNVs between genomes, CNVs overlapping by more than 50% in genomic location were considered shared between breeds. CNV related genes were defined on the basis of their sequence overlapping by at least 50% with a CNV.

### Phylogenetic tree construction

Phylogenetic trees were generated using SNPs detected from all breeds. First, reads that mapped to multiple genomic regions were filtered. Since Illumina-sequenced datasets possessed greater depth coverage than SOLiD-sequenced datasets, only regions that were covered by more than 10 uniquely mapped reads in all Illumina-sequenced data sets and more than 5 uniquely mapped reads in all SOLiD-sequenced datasets were used in this analysis. Further, only SNPs that fall within these regions and were present in at least one Illumina-sequenced breed and two SOLiD-sequenced breeds were considered for phylogenetic analysis. SNPs were concatenated to generate multiple sequence alignments, and phylogenetic trees generated by PhyML[61] with 1024 bootstrap replicates, using the Red Jungle Fowl as the outgroup. The final trees were visualized using FigTree[62].

### SNP calling of additional SOLiD sequenced datasets

Analyzing reads generated using the SOLiD platform required a different pipeline to detect SNPs. Bfast (v0.7.0)[63] converted fasta files and corresponding quality scores to csfastq files, which are normal outputs of the SOLiD platform and encode DNA sequences in colour space, the shrimp (v2.2.3)[64] aligner then mapped reads to the reference genome, with --max-alignments set to 5 to reduce reads that multi-mapped to many regions in the genome. Normal SNP calling steps were followed as before; Samtools and picard-tools were used again to sort, index, and remove duplicates from the alignment. Freebayes (v0.9.2) [65] was subsequently used to detect SNPs for SOLiD datasets with the flags – genotype-qualities –use-mapping-quality –genotyping-max-iterations 500.

### Functional annotation and analysis of genetic variants

SNPs were annotated with both ANNOVAR (v2016-02-01)[66] and Variant Effect Predictior (release 87)[67]. Variant Effect Predictor also provides information regarding the impact of variants on the basis of Sorting Tolerant From Intolerant (SIFT) score[68]. The vcf-isec package from VCFtools [69] was used to determine variants unique or shared across lines, for variants specific to broiler breeds, BCFtools[70] first merged all non-broiler variant files, and *vcf-isec* applied to compare broiler variants to the merged file to identify unique variants. SNP density for each gene was defined as the number of SNPs normalized by gene length, global SIFT score was defined as the average SIFT score of all variants in a gene normalized by gene length, and number of variants predicted to be deleterious for each gene. Ortholog[71], Gene Ontology[72], and functional annotations were retrieved from Ensembl Biomart[73] and Uniprot[74].

### Enrichment analysis

Genes with SNP densities in the 90^th^ percentile in each breed were subject to pathway enrichment analysis based on Gene Ontology (GO)[72] and Kyoto Encyclopedia of Genes and Genomes (KEGG)[75] categories. Only GO and KEGG categories containing at least 3 genes were considered. Enrichment analysis was performed by the R package clusterProfiler[76], and only categories with a p-value < 0.05 after Benjamini-Hochberg false discovery rate[77] correction were considered.

### Prioritization strategy

Based on SNP analyses described above, we defined five prioritization strategies to define transcripts of interest (**Figure 3**):

Strategy 1. Transcripts that have accumulated most SNPs in the last 50 years. Nonsynonymous SNP density increase between the 1957 and the 1978 broiler was compared to the nonsynonymous SNP density increase between the 1978 and Ross 308 broiler. For a transcript to be prioritized by this strategy, the difference in SNP density between the 1957 and 1978 broiler OR between the 1978 broiler and Ross 308 must be in the 95th percentile. Additionally, all SNP density changes between 1957 and 1978 and 1978 and Ross 308 must be positive (i.e. more SNPs in more recently bred lines)

Strategy 2. Transcripts with the most SNPs unique to fast-growing broilers. This strategy selected for transcripts with the highest number of variants (90th percentile after normalizing by transcript length) that belonged exclusively to the 1978 and/or Ross 308 subset OR the subset of SNPs exclusive only to Ross 308. The former subset (transcripts with high number of unique variants in 1978 and Ross 308 combined) includes transcripts that have not gained unique variants between 1978 and Ross 308 but have accumulated unique variants since the 1950s.

Strategy 3. Transcripts containing high impact variants (i.e. frameshift, start loss, and stop gain variants) only in Ross 308 or the 1978 broiler.

Strategy 4. Transcripts with decreasing SIFT score. For every transcript, the mean SIFT score is calculated, only transcripts that show a decrease in SIFT score when comparing the 1957, 1978, and Ross 308 broilers and are also in the bottom 50th percentile for either the 1978 or Ross 308 broiler are selected.

Strategy 5. Transcripts that have accumulated deleterious SNPs. Transcripts that show an increase in number of deleterious variants (SNPs with SIFT scores less than 0.05) when comparing the 1957, 1978, and Ross 308 broilers and possess the highest number of deleterious variants (99th percentile) after normalizing by transcript length.

## DECLARATIONS

Sequence data generated in this study is available through the sequence repository archive, hosted by the NCBI, under BioProject PRJNA561435

## ACKNOWLEDGEMENTS

This work was funded by grants from the Natural Sciences and Engineering Research Council of Canada (RGPIN-2014-06664 / RGPIN-2019-06852), Alberta Livestock and Meat Agency, Ontario Ministry of Agriculture, Food and Rural Affairs and the Canadian Poultry Research Council. High performance computing was provided by the Centre for Computational Medicine at the SickKids’ Research Institute.

