## Supplemental Table 1 for "Accumulation of genetic variants associated with immunity in the selective breeding of broilers"

**Supplemental Table 1. Breeds and Naming Convention for the 15 Samples Analyzed in this Study.**

The main text uses the abbreviation to refer to each breed. Genomes sequenced from pooled blood samples are specified in brackets in the “Name” column.

| **Study** | **Platform** | **Name** | **Abbreviation** |
| --- | --- | --- | --- |
| This study | Illumina NextSeq 500 | Ross 308 | Ross 308 |
|  |  | 1978 | 1978 |
|  |  | 1957 | 1957 |
| Fan *et al.*2013 (SRP022583) | Illumina Genome Analyzer II | Silkie | Silkie |
|  |  | Taiwan heritage L2 line | L2 |
| Rubin *et al*.*,*2010 (SRP001870) | SOLiD | Ross 308 (10 males) | CB1 |
|  |  | Undisclosed commercial broiler (10 females) | CB2 |
|  |  | Swedish layer (11 males) | WLA |
|  |  | Commercial White Leghorn (8 males) | WLB |
|  |  | High growth line (7 males, 4 females) | High |
|  |  | Low growth line (7 males, 4 females) | Low |
|  |  | Obese (10 males) | Obese |
|  |  | Rhode Island Red (8 males) | RIR |
|  |  | Swedish red jungle fowl (8 males) | RJFswe |
|  |  | Red jungle owl | RJF |
